# Lack of accelerated ovarian aging in a follicle-stimulating hormone receptor haploinsufficiency model

**DOI:** 10.1101/2022.10.07.511325

**Authors:** Kristen Mehalko, Minhoo Kim, Sanjana Paye, Kelly Koh, Ryan J. Lu, Bérénice A. Benayoun

## Abstract

Follicle-stimulation hormone (FSH) and FSH receptor (FSHR) signaling is essential for lifelong ovarian and endocrine functions in females. Previous studies have reported that *Fshr* haploinsufficiency in female mice led to accelerated ovarian aging, including anticipated progressive fertility decline, irregular estrus cycles, increased follicular atresia and premature ovarian failure at 7 to 9 months of age. Interestingly, these phenotypes resemble key characteristics of human menopause and thus *Fshr* haploinsufficiency was proposed as a promising research mouse model of menopause. However, the *Fshr* haploinsufficiency model had not been fully explored, especially at the molecular level. In this study, we characterized the ovarian and endocrine functions of a *Fshr* heterozygous knockout allele that was generated on the C57BL/6 genetic background as part of the Knockout Mouse Project (KOMP). Based on our analyses of these mice using a breeding assay, ovarian tissue histology and serum hormone quantifications (*i.e*. FSH, AMH, INHA) analyses, the KOMP *Fshr* heterozygous knockout female mice do not show the anticipated phenotypes of ovarian aging in terms of fertility and endocrine function. We further confirmed that the expression of *Fshr* is unaltered in the ovaries of the KOMP *Fshr* heterozygous knockout animals compared to wild-type. Together, our data suggests that the KOMP *Fshr* heterozygous knockout strain does not recapitulate the previously reported ovarian aging phenotypes associated to another model of *Fshr* haploinsufficiency.

## 1. Introduction

Mammalian female reproductive lifespan is limited by the fixed ovarian reserve predetermined at birth [1, 2]. In humans, ovarian aging is characterized by the progressive depletion of the ovarian reserve and deterioration of oocyte quality, culminating in menopause [1]. Age-associated loss of ovarian function not only leads to infertility, but is also accompanied by changes in circulating hormone levels, such as low-to-undetectable estradiol levels and high levels of follicle-stimulating hormone (FSH), luteinizing hormone (LH) and androgens [3]. Importantly, studies have shown that ovarian aging contributes to multisystem aging and frailty [4, 5]. Indeed, post-menopausal women are more susceptible to age-associated diseases, including neurodegeneration and osteoporosis [4, 6, 7]. Despite the significant health impact of menopause, the molecular mechanisms underlying the processes of ovarian aging are understudied, partly due to the limitations of current research models of menopause.

FSH is a pituitary gonadotropin which plays essential roles in the reproductive system [8, 9]. In females, FSH is required for stimulating growth and recruitment of follicles as well as survival of early antral follicles in the ovaries [10]. Elevated serum FSH level is a hallmark of menopause – due to the progressive decline in the number of recruited antral follicles, FSH levels gradually increase with age [11]. FSH functions by binding to the FSH receptor (FSHR), which is a specific G protein-coupled receptor present on ovarian granulosa cells [12, 13]. Studies have shown that the presence of FSHR is essential to FSH signaling and activity [9, 12]. High serum FSH levels in post-menopausal women have been correlated to low FSHR in the ovarian tissues [11]. Consistently, a decline in FSHR levels has been linked to ovarian hypo-stimulation and shown to lead to accelerated oocyte loss in mice [14]. Lack of FSHR was also shown to inhibit preovulatory and follicle development [9]. In contrast, high levels of FSHR have been found to slow reproductive aging and contribute to overall reproductive health [9].

*Fshr* haploinsufficiency has been described as a promising genetic mouse model of menopause [14]. Reminiscent of human menopause, *Fshr* heterozygous knockout female mice were shown to undergo anticipated progressive decline in fertility, estrus cycle irregularity and increased follicular atresia by early middle-age [14–16]. Additionally, *Fshr* haploinsufficient female mice were reported to undergo premature reproductive arrest, with endocrine profiles reminiscent of those observed in post-menopausal women (*i.e*. low levels of estradiol, and elevated levels of FSH, LH and androgens) [15, 16]. However, the potential of this strain as a mouse model of menopause has not been fully characterized at the molecular level.

Based on the previous findings on *Fshr* haploinsufficiency and its potential as a preclinical model for menopause-like states in laboratory mice, we decided to procure mice carrying an *Fshr* knock-out allele for further characterization. Specifically, we leveraged a *Fshr* heterozygous knockout mouse strain that had been generated by the Knockout Mouse Project (KOMP) and Mutant Mouse Resource and Research Center (MMRRC), hereafter referred to as *Fshr^+/−^*. After litter recovery, we started to characterize the ovarian health and aging phenotypes of *Fshr^+/−^* mice and their control littermates. Surprisingly, our results showed the KOMP *Fshr* heterozygous knockout female mice fail to recapitulate the previously characterized phenotypes observed with a different *Fshr* knockout allele. In this study, we describe the reproductive function of the KOMP *Fshr* heterozygous knockout female mice and discuss potential causes for the observed discrepancies in ovarian phenotypes.

## 2. Materials and methods

### 2.1. Husbandry

Mice were treated and housed in accordance with the Guide for Care and Use of Laboratory Animals. All experimental procedures were approved by the University of Southern California (USC)’s Institutional Animal Care and Use Committee (IACUC) and are in accordance with institutional and national guidelines. For externally procured animals, animals were allowed to acclimate in the animal facility at USC for 2-4 weeks prior to experimental use. The facility is on a 12-h light/dark cycle and animal housing rooms are maintained at 22°C and 30% humidity. For sample collection, all mice were euthanized via CO_2_ chamber asphyxiation succeeded by secondary cervical dislocation. Samples for this study were collected between 10am and 12pm. All animals were housed on approved IACUC protocol number 21155 at USC.

### 2.2. Generation of *Fshr* heterozygous knockout mice

Recovery of the *Fshr* heterozygous knockout mouse strain, C57BL/6N-*Fshr^tm1(KOMP)Vlcg^* (hereinafter referred to as “*Fshr*^+/−^”) was performed by the Jackson Laboratories (Bar Harbor, Maine). Briefly, cryo-archived sperm of C57BL/6N-*Fshr^tm1(KOMP)Vlcg^*/NjuMmucd (MMRRC, 047765-UCD) was purchased from MMRRC and used to fertilize eggs from C57BL/6J mice to recover *Fshr*^+/−^ animals [17]. The *Fshr^tm1(KOMP)Vlcg^* allele was previously generated as part of the Knockout Mouse Project (KOMP) [18]. The knockout allele was reported to host an insertion of the ZEN-Ub1 vector at genomic position 89,200,609 – 89,200,467 (GRCm38 assembly, Chromosome 17), thus leading to the interruption of the mouse *Fshr* locus, and a presumed null allele (Fig. S1A). We confirmed the position and nature of the invalidated allele (Fig. S1B-C).

The recovered litter was comprised of 2 *Fshr^+/−^* females, 1 *Fshr^+/−^* male and 3 wild-type females, and the mice were transferred to the animal facility at USC. Together with male C57BL/6J mice ordered from Jackson laboratories, these mice were used for breeding to expand the colony for study and phenotyping. After this first generation, mice were always bred with other mice from the colony to generate experimental animals.

Surprisingly, although the allele was reported to be generated in the C57BL/6 genetic background and the mice were recovered by fertilization of C57BL/6J eggs, this colony yields approximately 10-20% agouti coat-colored pups in addition to expected black coat-colored pups. Thus, we believe that it is unlikely that the revived *Fshr^+/−^* mice are of pure C57BL/6 background.

### 2.3. Genotyping and allele characterization

Genotypes of the mice were validated using PCR and PCR primers initially used by the Jackson Laboratories when recovering the *Fshr*^+/−^ mice (Table 1).

**Table 1.**
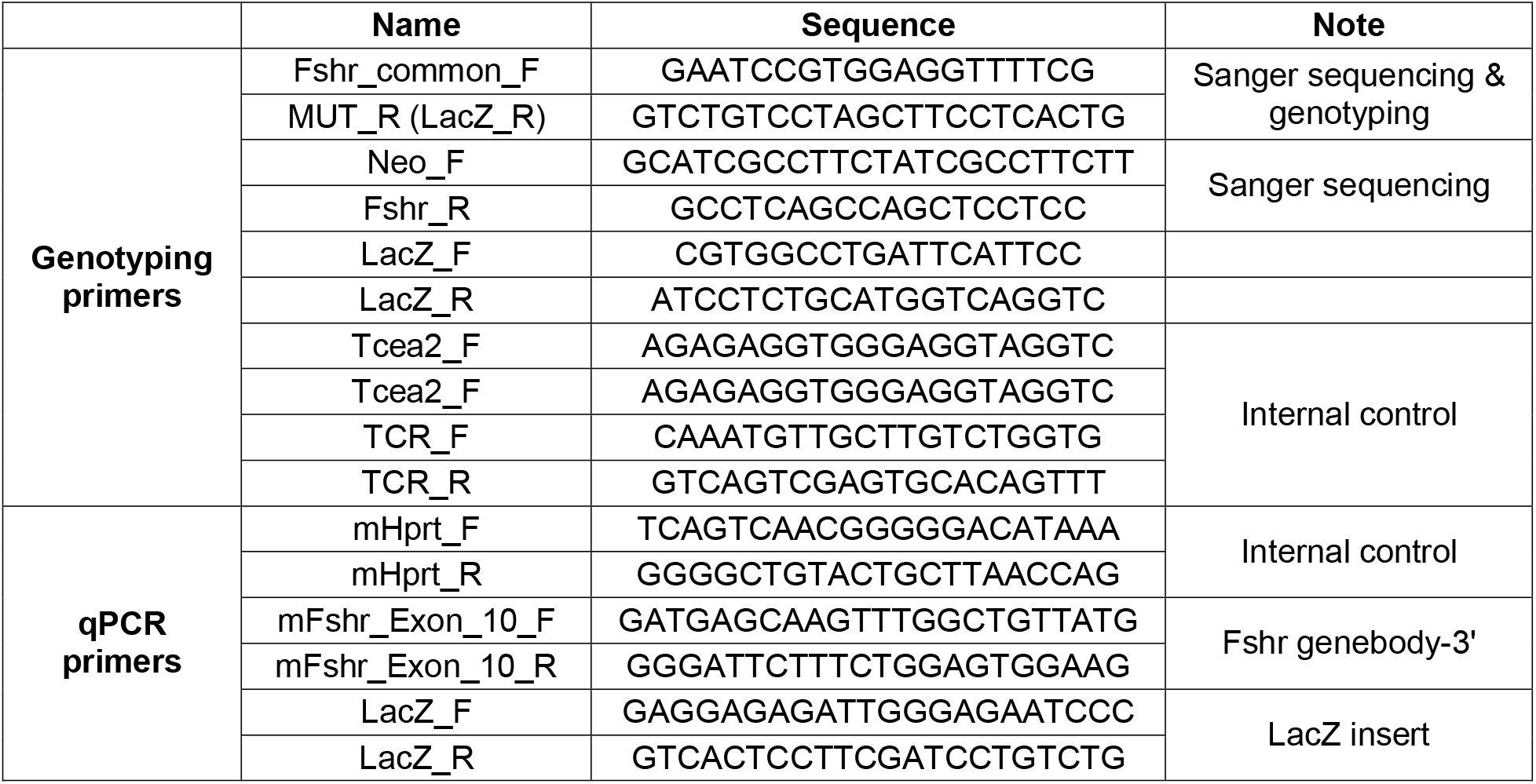
Primers used in the study.

Genomic DNA was prepared from tails using an in-house tail DNA lysis and extraction protocol (routine genotyping). Briefly, tail biopsies were suspended in tail lysis buffer (0.1M Tris-HCl pH 8.0, 5mM EDTA, 0.2% SDS and 0.2M NaCl) overnight and DNA was purified via isopropanol precipitation. DNA pellets were resuspended in 1X TE buffer. Alternatively, for higher yield and quality DNAA preparations, we also isolated genomic DNA from post-mortem liver tissue using the Monarch^®^ Genomic DNA Purification Kit (NEB, T3010S).

We also performed Sanger sequencing validation of the 5’- and 3’-end of the deletion cassette insertion site in the *Fshr* locus (Fig. S1B-C). Briefly, we used PCR to amplify a genomic fragment that spans −63bp and +132bp of the insertion start site and −380bp and +568bp of the insertion end site (see Table 1 for primers). After gel electrophoresis, the PCR products of 195bp and 948bp were gel purified using the NucleoSpin^®^ Gel and PCR Clean-up Kit (Machery-Nagel, 740609.250) and sent for Sanger sequencing at Azenta Life Sciences.

### 2.4. Fertility assay

Wild-type and *Fshr*^+/−^ females (n = 10 per group) that were born within a 20-day span were recruited and enrolled in the fertility assay (Table 2; Table S1). Age-matched C57BL/6J male mating partners were obtained from the Jackson Laboratory (Stock 000664) and allowed to acclimate in the animal facility at USC for 2-4 weeks prior to breeding. We initiated the breeding when females reached an average age of ~2.23 months (see Fig. S1E and Table 2 for specific ages), and paired them with 2-month-old male partners. Each dam was then allowed to birth four litters, with the exception of one wild-type dam (Mouse ID: 8), which did not produce four litters within the 19 weeks of the assay, likely due to an unusually long latency between her second and third litter (Table S1). Thus, mice reached about 6-7 months of age before the breeding assay was terminated. Litter numbers, number of pups per litter and latency between litters were recorded (Table S1). Importantly, data from the 3^rd^ litter from a *Fshr^+/−^* dam (Mouse ID: 11) and 4^th^ litter from a wildtype dam (Mouse ID: 8) are missing due to cannibalism and time constraints, respectively (Table S1).

**Table 2.** Metadata of mice enrolled in the study.

| Fertility assay |  |  |  |  |  |
| --- | --- | --- | --- | --- | --- |
| Mouse_ID | Genotype | Age at start of assay (months) | Mouse_ID | Genotype | Age at start of assay (months) |
| 1 | Fshr +/+ | 2.7 | 11 | Fshr +/- | 2.4 |
| 2 | Fshr +/+ | 2.6 | 12 | Fshr +/- | 2.4 |
| 3 | Fshr +/+ | 2.4 | 13 | Fshr +/- | 2.2 |
| 4 | Fshr +/+ | 2.3 | 14 | Fshr +/- | 2.2 |
| 5 | Fshr +/+ | 2.2 | 15 | Fshr +/- | 2.1 |
| 6 | Fshr +/+ | 2.2 | 16 | Fshr +/- | 2.1 |
| 7 | Fshr +/+ | 2.2 | 17 | Fshr +/- | 2.1 |
| 8 | Fshr +/+ | 2.1 | 18 | Fshr +/- | 2.1 |
| 9 | Fshr +/+ | 2.1 | 19 | Fshr +/- | 2.1 |
| 10 | Fshr +/+ | 2.1 | 20 | Fshr +/- | 2.0 |

| Ovarian histology & serum hormone quantification |  |  |  |  |  |
| --- | --- | --- | --- | --- | --- |
| Mouse_ID | Genotype | Age at euthanasia (months) | Mouse_ID | Genotype | Age at euthanasia (months) |
| 21 | Fshr +/+ | 4 | 29 | Fshr +/- | 4 |
| 22 | Fshr +/+ | 4 | 30 | Fshr +/- | 4 |
| 23 | Fshr +/+ | 4 | 31 | Fshr +/- | 4 |
| 24 | Fshr +/+ | 4 | 32 | Fshr +/- | 7 |
| 25 | Fshr +/+ | 7 | 33 | Fshr +/- | 8 |
| 26 | Fshr +/+ | 9 | 34 | Fshr +/- | 8 |
| 27 | Fshr +/+ | 9 | 35 | Fshr +/- | 9 |
| 28 | Fshr +/+ | 9 |  |  |  |

### 2.5. Serum hormone level quantification and ovarian histology analysis

Young (4-months) and middle-aged (7-9-months) virgin wild-type (n = 4 young and n = 4 middle-aged) and *Fshr*^+/−^ females (n = 3 young and n = 4 middle-aged) were used to collect ovaries for histological analysis and serum for reproductive hormone level quantification (Table 2).

For each animal, we collected one ovary for RNA extraction, which was flash-frozen in liquid nitrogen upon harvest and kept at −80°C until processing (see below). The contralateral ovary was fixed in Bouin’s solution (Sigma, HT10132) for 24 hours at room temperature and transferred to 70% ethanol for storage. Paraffin embedding, tissue sectioning and staining with Hematoxylin and Eosin (H&E) were performed by the USC Norris Comprehensive Cancer Center Translational Pathology Core Facility. H&E staining slides were imaged on the Echo Revolve microscope platform. Two sections per ovary were used to count the number of primordial, primary, secondary, and antral follicles and corpus luteum by three blinded observers. Median counts across the 3 observers were used for data analysis (Fig. 2C). The mean of follicle counts from the same experimental groups (*i.e*. Young wild-type, young Fshr^+/−^, middle-aged wild-type and middle-aged Fshr^+/−^) was used to generate the stacked bar chart in Fig. 2C.

After euthanasia, blood was collected directly from the heart. Blood was allowed to clot at room temperature for an hour, and serum was separated using MiniCollect^®^ Serum Tube (Greiner, 450472), then snap frozen in liquid nitrogen. Quantification of serum FSH (Millipore Pituitary Panel Multiplex kit), AMH (The Rat and Mouse Anti-Müllerian hormone (AMH) enzyme-linked immunosorbent assay (ELISA) kit, Ansh Labs, AL-113) and INHA (The animal Inhibin A enzyme linked immunosorbent assay (ELISA) kit, Ansh Labs, AL-161) levels were performed by the University of Virginia Center for Research in Reproduction Ligand Assay and Analysis Core.

### 2.6. RT-qPCR quantification of ovarian *Fshr* mRNA expression levels

Flash-frozen ovaries collected from young and middle-aged virgin wild-type and *Fshr*^+/−^ females (see above) were used for RNA extraction, cDNA synthesis and quantitative PCR(qPCR). For RNA extraction, ovarian tissues were transferred into Lysing Matrix D tubes (MP, 6913500) together with 200μL of TRIzol^®^ Reagent (Ambion, 15596018). The tissue was homogenized using a BeadBug 6 microtube homogenizer for 3 rounds of 30 seconds at 3000 rpm (Benchmark Scientific, 422V16). RNA was purified from homogenized tissue using the Direct-Zol RNA Miniprep kit (Zymo Research, R2052), according to the manufacturer’s instructions. cDNA was synthesized using the Maxima H Minus First Strand cDNA Synthesis Kit (Thermo Scientific, K1652) and RT-qPCR quantification were performed using the MIC Tubes and Caps kit (Bio Molecular Sytems, 71-101) on the Magnetic Induction Cycler (MIC) machine (Bio Molecular Systems, MIC-2). micPCR v2.12.6 was used to capture and quantify Ct values. Primers and sequences used for qPCR reactions can be found in Table 1.

### 2.7. Code and data availability

The raw data and R scripts for data plotting and statistics are available at https://github.com/BenayounLaboratory/Fshr_mouse_characterization.

## 3. Results

### 3.1. KOMP *Fshr^+/−^* female mice show normal fertility

We obtained an *Fshr* heterozygous knockout mouse strain (hereinafter referred to as *Fshr*^+/−^) from the Knockout Mouse Project (KOMP) and Mutant Mouse Resource and Research Center (MMRRC) [17, 18]. Specifically, cryo-preserved sperm of the C57BL/6N-*Fshr^tm1(KOMP)Vlcg^* strain, generated as part of the KOMP using the Velocigene targeted allele method (Fig. S1A) [19], was obtained from MMRRC and revived using oocytes from C57BL/6J mice at the Jackson Laboratories. Insertion of the deletion cassette in the *Fshr* locus and genotypes of the animals were confirmed via Sanger sequencing and PCR, respectively (Fig. S1B-D, S2). Due to our interests in understanding the impact of *Fshr* haploinsufficiency on ovarian function, our assays focused only on female animals from the colony.

Surprisingly, although the cryo-preserved sperm for *Fshr*^+/−^ mice was reported to have been generated on the C57BL/6N background and the egg donors for the recovery were from the C57BL/6J background, we observed agouti color-coated pups in our colony (approximately 10-20%), including in the recovered litter. Previously, subfertility in independent *Fshr*^+/−^ mouse strains was reported by two independent groups [14, 20]. To check any changes in the fertility of our *Fshr*^+/−^ mice, 20 female mice were enrolled in a fertility assay (n = 10 per genotype, wild-type vs. *Fshr*^+/−^), paired with C57BL/6J wild-type male mice (Fig. 1A, S1E; Table S1). In contrast to previous reports, we did not observe any significant differences in litter size, litter number or latency between litters between wild-type and *Fshr*^+/−^ mice (Fig. 1B-E; Table S1). Additionally, average litter size was comparable for mothers of different ages (~3 *vs*. ~7 months, *i.e*. litter 1 *vs*. litter 4), regardless of genotype (Fig. 1B-C). Furthermore, neither wild-type nor *Fshr*^+/−^ mice showed age-related decline in litter size by 7 months of age (Fig. 1B-C). This contrasts with the work by Danilovich and Sairam, where *Fshr*^+/−^ dams showed a two-fold reduction in reproductive success and litter size at 3, 7 and 12 months of age [14]. Importantly, our observed average litter sizes, regardless of dam genotype, are consistent with reported average litter sizes of 5.6 for C57BL/6 mice or 4.9 for SV129 mice [21, 22]. Overall, our data shows that the KOMP *Fshr*^+/−^ female mice do not show any accelerated age-related decline in fertility compared to wild-type female littermates.

**Fig. 1.**
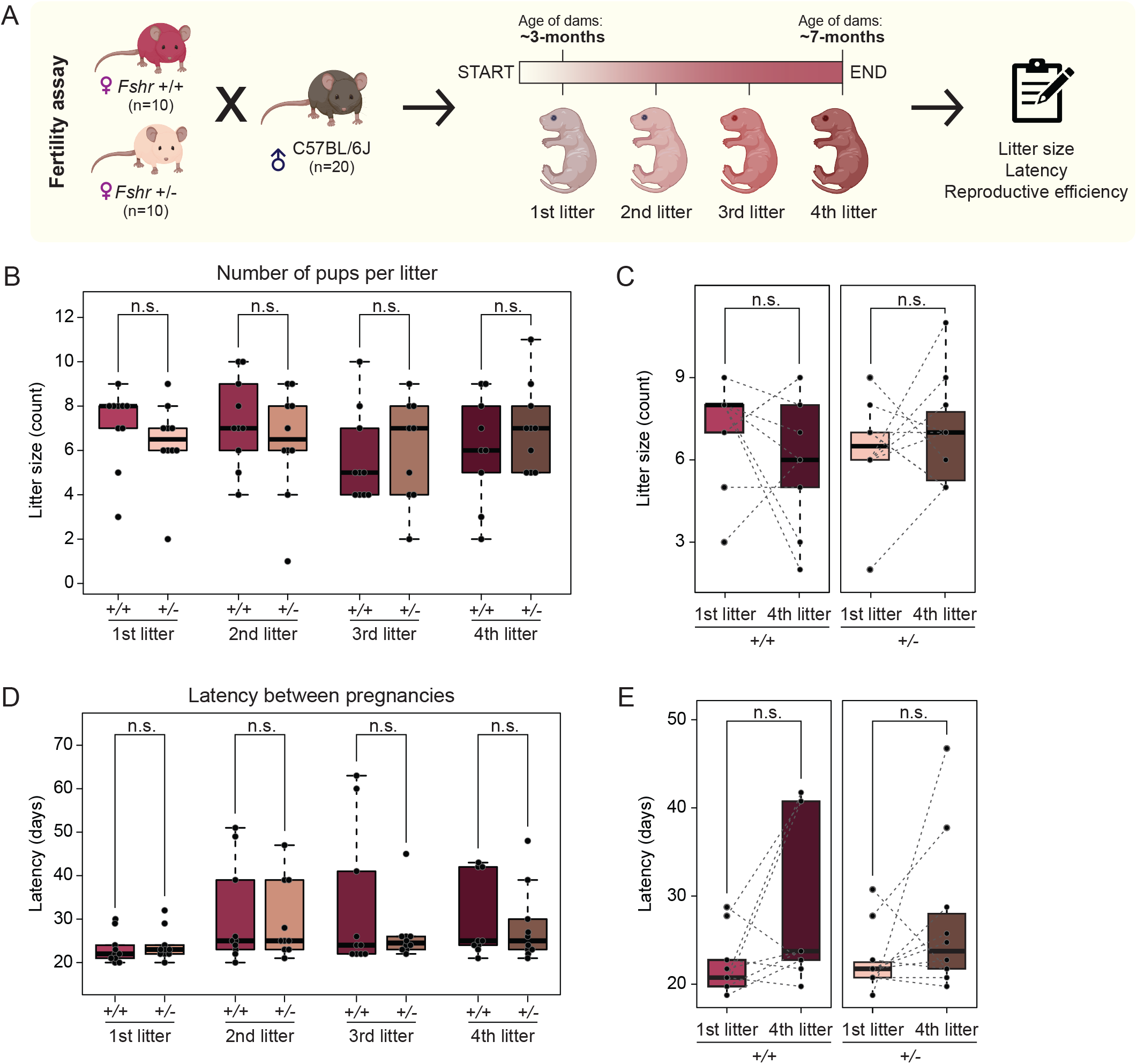
Fertility assay of wild-type and *Fshr^+/−^* female mice show a lack of significant genotype- or age-related difference in litter sizes and latency periods. (A) Schematic diagram of the fertility assay and mice enrolled. Panel was made by BioRender.com. (B) Number of pups birthed by each dam for each litter 1-4 (n = 9-10 per group). Data from a *Fshr^+/−^* dam that cannibalized her 3^rd^ litter before data recording was omitted from quantification since it could not be precisely recorded (refer to Table S1, Mouse ID: 11). (C) Paired comparison of number of pups in 1^st^ and 4^th^ litters at approximately 3 and 7 months of age, respectively, between wild-type and *Fshr^+/−^* dams (n = 10 per group). Data is replotted from (A) for ease of visual comparison. (D) Latency period between birth of each litter of each dam (n = 9-10 per group). (E) Paired comparison of latency periods for 1^st^ and 4^th^ litters between wild-type and *Fshr^+/−^* dams (n = 9-10 per group). Data is replotted from (D) for ease of visual comparison. For (D) and (E), due to time constraints, a wild-type dam was euthanized before birthing the 4^th^ litter and thus was omitted from quantification (refer to Table S1, Mouse ID: 8). Significance in non-parametric two-sided Wilcoxon rank-sum tests are reported for panels. +/+: *Fshr*^+/+^. −/−: *Fshr*^−/−^. n.s.: Not significant.

### 3.2. *Fshr*^+/−^ female mice show normal ovarian histology and serum FSH, AMH and INHA levels

It is well documented that FSH and FSHR signaling is important for the regulation of ovarian follicle maturation [13, 23]. Additionally, studies have suggested a protective role of FSH and FSHR signaling in the ovaries through inhibition of follicular atresia [24, 25]. Indeed, Danilovich and Sairam found that ovaries of *Fshr*^+/−^ female mice showed significant oocyte loss and follicular atresia by 7-months of age [14]. The same study found that, by 7-months of age, *Fshr*^+/−^ ovaries showed a ~75% decrease in total follicle numbers compared to 3-months of age [14].

To assess any differences in the ovarian landscape in our *Fshr*^+/−^ mice compared to wild-type littermate controls, we collected ovaries from young (4-months) and middle-aged (7-9-months), wild-type and *Fshr*^+/−^ female mice for histology analysis (Fig. 2A). Quantification of follicles revealed little difference in follicle counts between wild-type and *Fshr*^+/−^ mice, regardless of age (Fig. 2B-C).

**Fig. 2.**
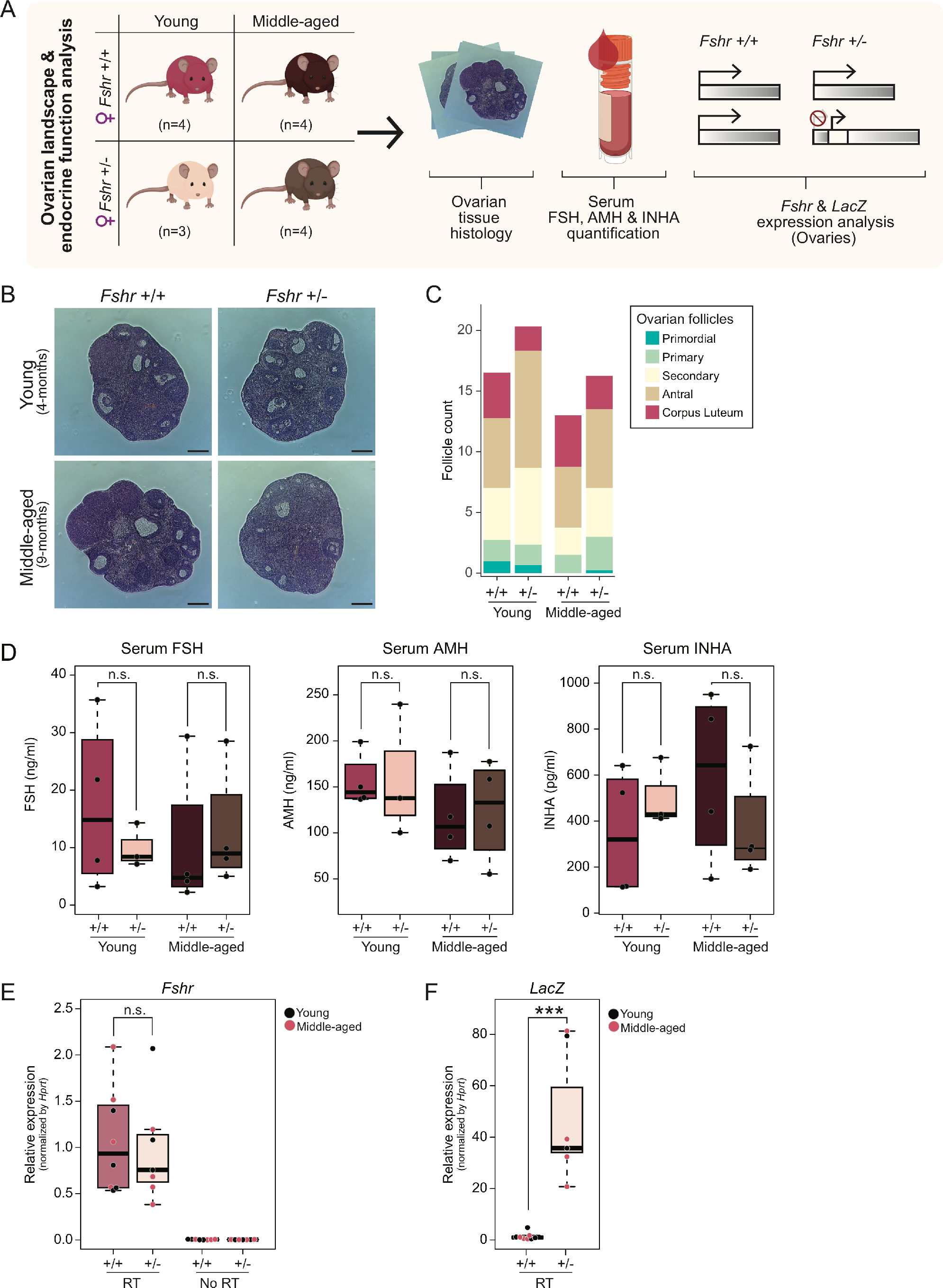
Lack of accelerated ovarian aging phenotypes in KOMP *Fshr^+/−^* females. (A) Schematic diagram of ovarian landscape and endocrine function assessment. (B) Representative images of ovarian tissue H&E staining from wild-type *vs. Fshr^+/−^* at youth *vs*. middle-age. Scale bar = 300μm. (C) Quantification of follicle counts from ovarian tissue H&E staining data (2 slides per ovary, counted by 3 blinded observers). (D) Boxplots of serum FSH, AMH and INHA quantification data. (E and F) Relative expression of *Fshr* gene body (p ~ 0.87) (E) and the *LacZ* reporter (p ~ 3.1×10^−4^) (F). Significance in non-parametric two-sided Wilcoxon rank-sum tests reported for all relevant panels. +/+: *Fshr*^+/+^. −/−: *Fshr*^−/−^. n.s.: Not significant.

In parallel, we decided to quantify endocrine markers of ovarian health and reserve in the serum of these mice (*i.e*. FSH, Anti-Müllerian Hormone (AMH), Inhibin A (INHA)) (Fig. 2D). Although a work by Dierich et al. showed minimal differences in the FSH levels in the *Fshr*^+/−^ female mice compared to wild-type controls, the study used relatively young animals (3-month-olds) for quantification [20]. We hypothesized that changes in FSH levels (specifically, an increase in FSH levels) in the *Fshr*^+/−^ may be detectable in older animals (*e.g*. 7-9-months), at the age at which premature ovarian failure had previously been described for *Fshr* haploinsufficient mice. Indeed, FSH levels are widely used as an indicator of ovarian aging, with sustained high FSH levels being used as a diagnostic marker for menopause and premature ovarian failure in humans [26]. INHA is a gonadal glycoprotein produced by granulosa cells, which is required for suppression of FSH secretion [27]. Importantly, low INHA levels are a marker of reduced ovarian function [28]. Lastly, AMH is a glycoprotein hormone that is produced by ovarian granulosa cells and is used as a marker of the ovarian reserve in the clinic, with decreased AMH levels being a marker of reduced follicle reserve in the aging ovary [29–31]. Consistent with our fertility assay results, serum hormone quantification showed similar levels of FSH, INHA and AMH levels between wild-type and *Fshr*^+/−^ animals, regardless of age (Fig. 2D). Collectively, our data indicates that the KOMP *Fshr*^+/−^ female mice have normal ovarian function compared to the wild-type controls and thus lack the previously described accelerated ovarian aging phenotype.

### 3.3. Normal expression of *Fshr* in the ovaries from KOMP *Fshr*^+/−^ mice

Accelerated ovarian aging phenotypes that had been described in mice carrying other alleles for *Fshr*^+/−^ was expected to result from decreased *Fshr* mRNA expression levels in relevant tissues (*e.g*. ovary), and thus decreased protein levels leading to decreased signaling through the Fsh/Fshr pathway. Reduced FSH signaling could then lead to premature depletion of the ovarian reserve, and thus accelerated ovarian aging. Due to the lack of ovarian phenotypes in the KOMP *Fshr*^+/−^ mouse, we surmised that the wild-type *Fshr* allele may be able to compensate expression levels of *Fshr* in our strain, and thus prevent the occurrence of observed premature ovarian aging phenotypes. We used RT-qPCR to measure expression levels of *Fshr* and *LacZ* (*i.e*. deletion cassette reporter) in the ovarian tissues from wild-type and *Fshr*^+/−^ animals (Fig. 2E-F). Consistent with the lack of observed ovarian defects, we did not observe a decrease in the *Fshr* expression in the *Fshr*^+/−^ mice ovaries compared to wild-type controls (Fig. 2E). As an important control, expression of *LacZ* was robustly observed only in the ovaries from *Fshr*^+/−^ mice, consistent with the presence of the invalidated allele (Fig. 2F). Together, our results are consistent with the notion that the KOMP *Fshr* knockout allele does not lead to reduced *Fshr* expression levels in key tissues, with expression of the wild-type allele compensating for the invalidated allele.

## 4. Discussion

In this study, we investigated the ovarian health and endocrine function of the *Fshr* heterozygous knockout mouse strain that was developed as part of the Knock-Out Mouse Project (KOMP) [18]. We mapped the precise genomic site of insertion of the deletion cassette (ZEN-Ub1) in the mouse *Fshr* locus, as well as the expression of *Fshr* and the *LacZ* reporter in the ovaries of *Fshr*^+/−^ animals. However, we could not observe changes in the ovarian *Fshr* expression in the KOMP *Fshr*^+/−^ female mice compared to wild-type controls. Consistently, our fertility assay, ovarian tissue histology and serum hormone quantification (FSH, AMH and INHA) analyses were consistent with normal (rather than the expected accelerated) ovarian aging in the KOMP *Fshr*^+/−^ female mice.

Notably, we designed our fertility assays to minimize the numbers of required dams, and to be able to follow the reproductive performance of each dam longitudinally with aging. In contrast, in the work by Danilovich and Sairam, virgin females (both wild-type and *Fshr*^+/−^) were enrolled in the fertility assay at different time points (3, 7 and 12 months of age) [14]. Although this difference in setup may impact observation for dams at age 7 months, Danilovich and Sairam reported significant decreases in litter sizes as early as 3 months of age in *Fshr*^+/−^ females compared to age-matched wild-type controls, with differences increasing with age [14]. In our data, KOMP *Fshr*^+/−^ females had normal litter sizes at 3 months of age (directly comparable to [14]), and normal subsequent litter sizes, number and latency up to 7 months of age (Fig. 1B-E). Since trajectories of reproductive function between virgins and female mice that have been continuously bred are comparable [32], the differences in breeding assay setup are unlikely to explain the lack of fertility impairment in the KOMP *Fshr*^+/−^ mice. Typically, C57BL/6 female mice are fertile between 2 and 10 months of age, and start to experience fertility decline at ~7-8 months [33]. We enrolled ~2-month-old mice and longitudinally tracked their reproductive function in order to obtain a comprehensive overview of the impact of *Fshr* haploinsufficiency on reproductive function.

Importantly, our ovarian histology was performed in virgin animals, as in Danilovich and Sairam [14]. Although they observed clear changes in ovarian morphology and follicle content at 3 and 7 months of age in *Fshr*^+/−^ animals, we did not observe substantial changes in ovarian histology in comparable conditions for the KOMP *Fshr*^+/−^ animals (Fig. 2B-C). Overall, our observations using the KOMP *Fshr*^+/−^ mice contrast with previous observations on other *Fshr* haploinsufficient mouse models, which showed reduced reproductive performance in both the C57BL/6 and SV129 backgrounds [14, 20]. Below, we discuss potential contributing factors that may explain the discrepancies between previous reports using other *Fshr* invalidation alleles and our observations using the KOMP *Fshr* invalidation allele.

### Differences in gene knockout methods

In previous studies by Danilovich and Sairam, and Dierich et al., *Fshr* invalidation was induced by a 11kb deletion of the *Fshr* locus that encompasses a portion of exon I and intron I [14]. On the other hand, the KOMP *Fshr* knockout strain was generated using the Velocigene targeted allele method in which only a short ~143bp fragment following the start codon of the *Fshr* gene was replaced by a ~6kb cassette (ZEN-Ub1) (Fig. S1A) [19]. Although a 6kb insertion of exogenous DNA in the *Fshr* locus is expected to result in a non-functional allele, the specific knockout method used to generate our *Fshr*^+/−^ strain may have contributed to genetic compensation of *Fshr* expression [34]. For example, a recent study showed that the expression and degradation of mutant mRNA are required to induce genetic compensation by the wild-type allele in both mouse and zebrafish through a sequence-dependent mechanism [35]. Hence, it is possible that the specific mutant mRNA expressed and degraded in the tissues of the KOMP *Fshr*^+/−^ mice may have promoted genetic compensation and upregulation of the wild-type allele, whereas the mutant mRNA in the previous knockout alleles did not.

### Differences in mouse genetic background

The two key studies that characterized the impact of *Fshr* haploinsufficiency on female reproductive function used pure genetic mouse backgrounds, C57BL/6 and SV129, to evaluate the impact of *Fshr* haploinsufficiency [14, 20]. Our study was designed to use a pure C57BL/6 background, based on the generation of the KOMP *Fshr^tm1(KOMP)Vlcg^* allele on the C57BL/6N background, and the use of the Jackson laboratories C57BL/6J oocytes for recovery of the strain. However, starting from the recovered litter that we obtained, we always observed a mix of black and agouti coat-colored pups, supporting the notion that the allele was generated in mixed background mice (although the specific background cannot be easily elucidated based on the information that we have from KOMP).

It is well documented that genetic background can have significant effects on expression and penetrance of phenotype [36]. For example, knockout of the adenine phosphoribosyltransferase (*Aprt*) gene showed earlier onset of illness (3-4 weeks vs. 3-4 months) and shorter lifespan (75 days *vs*. 180 days) in C57BL/6J mice compared to Black Swiss mice [37]. Additionally, studies have investigated the effects of pure and mixed backgrounds on phenotypes of single-locus mutations [38]. For example, most TGFβ1 null mutants generated in mixed backgrounds, such as SV129 x C57BL/6, showed a survival rate of 50%, whereas only 1% of those in pure C57BL/6 inbred background survived to birth [38]. Given that we observed mixed coat colors in our colonies, the mixed genetic background may have led to the protection of mouse ovaries against the impact of *Fshr* haploinsufficiency.

Together, our observations suggest that the KOMP *Fshr* Velocigene targeted allele should not be used as a heterozygote genotype to elicit accelerated ovarian aging phenotypes, likely due to multiple factors leading to dosage compensation. Importantly, although such dosage compensation is not expected to be an issue for the study of full knockout phenotypes (*i.e*. in mice carrying two targeted alleles), this precludes the use of this strain to study *Fshr* haploinsufficiency-driven accelerated ovarian aging.

## Acknowledgments

Some panels were made with BioRender.com.

We thank Chan Boriboun for initial help on propagation and maintenance of the *Fshr* mouse strain at USC. We also would like to thank Dr. Marc Vermulst for his generosity in letting us use the Echo Revolve microscope for ovarian tissue histology imaging. This work was supported by GCRLE-2020 post-doctoral fellowship from the Global Consortium for Reproductive Longevity and Equality at the Buck Institute, made possible by the Bia-Echo Foundation to M.K., NIA T32 AG052374 predoctoral fellowship and a Diana Jacobs Kalman/AFAR Scholarships for Research in the Biology of Aging to R.J.L., and GCRLE-0520 Junior Scholar Award from the Global Consortium for Reproductive Longevity and Equality at the Buck Institute, made possible by the Bia-Echo Foundation to B.A.B.

The mouse strain used for this research project was created by the Mouse Biology Program (www.mousebiology.org) at the University of California Davis using vector 047765-UCD, generated by the trans-NIH Knock-Out Mouse Project (KOMP) and obtained from the MMRRC (www.mmrrc.org). NIH grants to Velocigene at Regeneron Inc (U01HG004085) and the CSD Consortium (U01HG004080) funded the generation of gene-targeted vectors and ES cells for over 8,500 genes for the KOMP Program. Serum hormone quantification was performed by the University of Virginia Center for Research in Reproduction Ligand Assay and Analysis Core (supported by the Eunice Kennedy Shriver NICHD/NIH Grant R24HD102061). Ovarian histological analysis was performed by the Translational Pathology Core at the USC Norris Comprehensive Cancer Center (supported by NCI P30 CA014089).

## Author Contributions

B.A.B. conceived the project. K.M. and R.J.L. expanded and maintained the mouse colonies. K.M., S.P., K.K. and M.K. performed the genotyping PCR reactions. M.K. performed the ovarian histology, serum hormone quantification and RT-qPCR analyses. K.M., S.P., M.K. and B.A.B. wrote the manuscript with input from all authors. All authors edited and commented on the manuscript.

## Legend to Supplementary Figures

**Fig. S1. Sanger sequencing of 5’- and 3’-end of the *Fshr* KO deletion cassette insertion site and genotyping PCR agarose gel electrophoresis data**.

(A) Schematic diagram of *Fshr* wild-type and knockout alleles (GRCm38 assembly). Diagrams are not drawn to scale. (B and C) Sanger sequencing chromatogram of 5’- and 3’-end of *Fshr* KO deletion cassette (ZEN-Ub1) insertion site. (D) Genotyping PCR agarose gel electrophoresis data of all animals used in this study. (E) Age of dams (*Fshr^+/+^* vs. *Fshr^+/−^*) at initiation of fertility assay. Significance in non-parametric two-sided Wilcoxon rank-sum tests reported for all relevant panels (p ~ 0.26). +/+: *Fshr*^+/+^. −/−: *Fshr*^−/−^. n.s.: Not significant.

**Fig. S2. Original electrophoresis agarose gel images**.

Uncropped agarose gel images corresponding to genotyping PCR gel electrophoresis data in Fig. S1D.

## Inventory of Supplementary Table

**Table S1. Raw data of fertility assay**.

