## Supplementary figures and images for "Lack of accelerated ovarian aging in a follicle-stimulating hormone receptor haploinsufficiency model"

### Fig. S1

#### Supplementary Figure 1

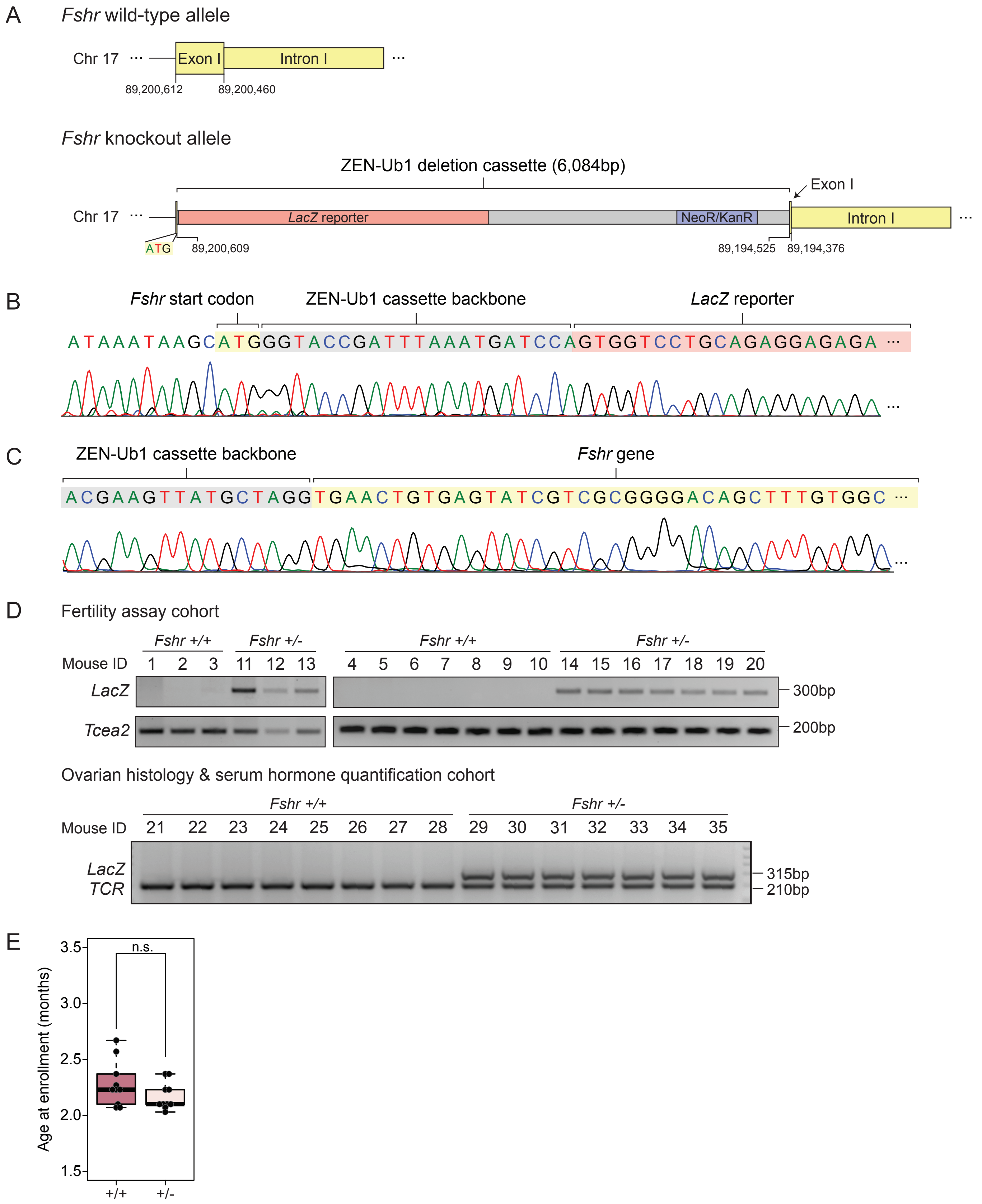
