## Supplementary material for "Lack of accelerated ovarian aging in a follicle-stimulating hormone receptor haploinsufficiency model": Fig. S2

Supplementary Figure 2

A

Fertility assay cohort

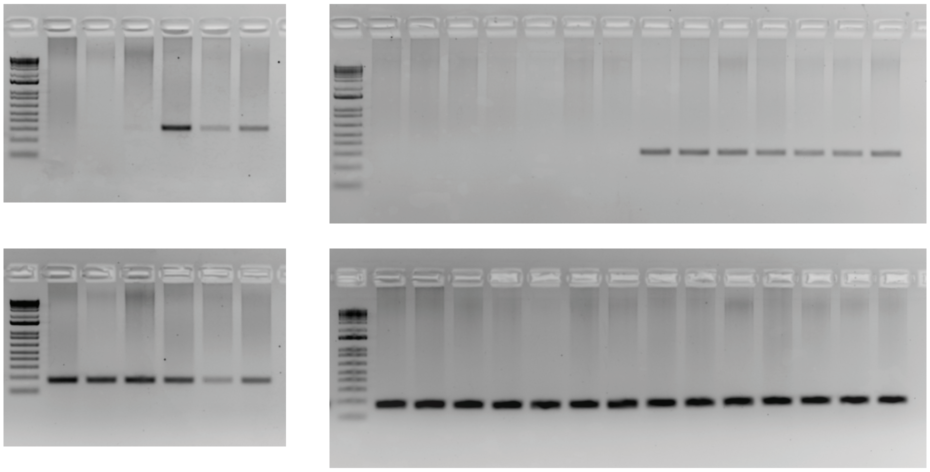

Ovarian histology & serum hormone quantification cohort

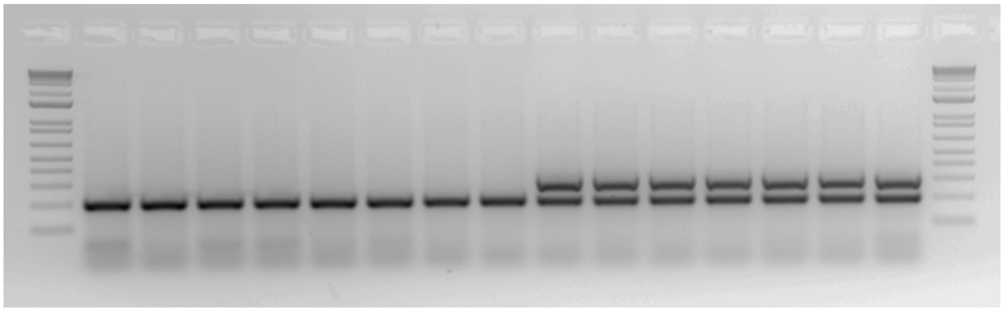
